# Computational profiling of hiPSC-derived heart organoids reveals chamber defects associated with Ebstein’s anomaly

**DOI:** 10.1101/2020.12.24.424346

**Authors:** Wei Feng, Hannah Schriever, Shan Jiang, Abha Bais, Dennis Kostka, Guang Li

## Abstract

Heart organoids have the potential to generate primary heart-like anatomical structures and hold great promise as in vitro models for cardiac disease. However, their properties have not yet been carefully studied, which hinders a wider spread application. Here we report the development of differentiation systems for ventricular and atrial heart organoids, enabling the study of heart disease with chamber defects. We show that our systems generate organoids comprising of major cardiac cell types, and we used single cell RNA sequencing together with sample multiplexing to characterize the cells we generate. To that end, we also developed a machine learning label transfer approach lever-aging cell type, chamber, and laterality annotations available for primary human fetal heart cells. We then used this model to analyze organoid cells from an isogeneic line carrying an Ebstein’s anomaly associated genetic variant, and we successfully recapitulated the disease’s atrialized ventricular defects. In summary, we have established a workflow integrating heart organoids and computational analysis to model heart development in normal and disease states.

## Introduction

Human induced pluripotent stem cells (hiPSCs) have been shown to differentiate into beating heart muscle cells (cardiomyocytes, CMs) with monolayer differentiation protocols, and into heart organoids comprised of a variety of cell types with three-dimensional differentiation systems.^1,2 3,4^ While monolayer differentiation protocols are able to produce very pure populations of cells, they are not able to model the three-dimensional spatial microenvironments of cardiac development; therefore these protocols may not be appropriate to study congenital heart defects (CHDs), the most frequently observed type of malformation at birth and the most common cause of infant death due to birth defects in the United States^5^. Organoids, on the other hand, are generated using three-dimensional differentiation methods, which enables them to develop anatomical context through self-assembly. This has already been leveraged to study developmental processes in several tissue and organ systems like brain, intestine, and kidney.^6–8^ Also in the context of heart development several three-dimensional differentiation protocols have been published,^2–4^ but their chamber identities have not been carefully investigated and applied to study CHD. Therefore, we established two three-dimensional differentiation protocols geared towards producing atrial and ventricular heart organoids, respectively. This approach then allowed us to study chamber defects in the context of CHDs in general and for Ebstein’s anomaly (EA) in particular.

EA as a rare but serious CHD occurs in about 1 per 200,000 live births and accounts for less than 1% of all cases of CHDs^9^. EA patients suffer from heart chamber malformations, including enlarged right atrium (RA), reduced right ventricle (RV), and abnormal tricuspid valves. Genetic causes play a role in EA, albeit the disease is genetically heterogeneous. Known genetic causes include chromosomal alterations (like copy number variations) and single gene defects in cellular structural proteins, signaling molecules and cardiac transcription factors.^10^ Specifically, multiple sequence variations within the homeobox-containing cardiac transcription factor *NKX2-5* have been associated with EA.^10,11^ Given that *NKX2-5* is a transcription factor with key roles in cardiac development^12–15^, and because knock out experiments in zebrafish and mouse have demonstrated that *NKX2-5* is involved in chamber specification in the developing vertebrate hearts,^16,17^ we were interested to further investigate a specific EA-associated variant in the coding sequence of *NKX2-5* converting a cytosine to an adenine (c.673C>A).^11^

Single cell RNA sequencing (scRNA-seq) enables the study of transcriptional profiles of individual cells, and it has successfully been used to study and elucidate disease etiology for CHD.^18–20^ Commercial droplet-based methods (like the 10X genomics platform) have been shown to capture a large diversity of cell types, and they can be used to assay a large number of cells in each experiment.^21^ We used this approach to characterize organoids generated by our protocols at different differentiation time points, and to compare wild type heart organoids with organoids that were genetically modified to carry the *NKX2-5* c.673C>A variant. A major consideration in the design of scRNA-seq experiments are batch effects, which arise when samples are processed in separate groups and have the potential to severely confound analysis results and downstream conclusions.^21,22^ Therefore we used the MULTI-seq approach that (through lipid-based sample barcoding) enables multiplexing of different samples for library preparation and sequencing.^23^

For comprehensive molecular characterization of cardiac cells based on scRNA-seq, we used machine learning to implement a label transfer approach (based on random forests) that allowed us to leverage information about cell type, heart chamber (atrial vs. ventricular) and laterality (left vs. right side) available for primary human fetal cells^18,24,25^ in the context of our in vitro system. The random forest learning algorithm is a machine learning method that has been successfully employed in the context of scRNA-seq data annotation,^17^ and we adopted and modified this approach to generalize well across different sequencing platforms, and to include an anomaly detection step to highlight cells that are likely not heart-related. This enabled us to characterize the differentiation protocols we established and to compare wild type to genetically modified cells.

Overall, we find that our differentiation approach generated organoids with atrial and ventricular heart cells. Single cell transcriptional profiling in combination with the label transfer approach we developed was able to identify a range of cardiac cells in our organoids. Comparison of cells from wild type organoids with cells from organoids with the EA-associated genetic lesion c.673C>A identified chamber developmental defects. Additionally, we found genes down-regulated in mutant cells are related to striated muscle differentiation, while up-regulated genes are related to energy and metabolism, illustrating specific molecular consequences of this genetic manipulation in the context of heart development. This finding suggests that our overall approach is a promising option for characterizing lineage defects and the functional roles of genetic variants in CHDs in general.

## Results

### Generation of ventricular heart organoids

In order to generate ventricular heart organoids, we established a three-dimensional differentiation protocol by sequentially modulating the *WNT* signaling pathway, which is largely similar to the established monolayer differentiation protocols.^2,26^ This allowed us to differentiate two hiPSC lines (WTC line with ACTN2-eGFP reporter and SCVI114 line) into cardiac lineages in organoid (Org) and monolayer (ML) systems (**Figure 1A**). Beating cells and ACTN2-eGFP signal were observable at day 15 and 30 in both protocols (**Figure 1B**, **Video 1**). Interestingly, slightly lower beating rates and shorter sarcomere lengths were observed in organoid cells compared to monolayer cells (**Figure S1A**, **C**). Organoid size increased throughout early stages of differentiation and remained stable between day 15 and 30, but the variance increased markedly after day 7 (**Figure 1C**) when the cells had been transferred from AggreWell to 6-well plates. Transverse sectioning of the organoids revealed varied internal structures, which we grouped into three categories: intact, holes, and cavities, with the latter being the largest group of organoids we observed (**Figure 1D**). Staining with cardiac troponin T (cTNT), a marker for cardiomyocytes, we found the majority of organoids with cavities (90.9%) and holes (71.4%) stained cTNT positive, while most organoids with intact structures (75%) were cTNT negative (**Figure S1D**). Quantification of cTNT-positive areas confirmed that organoids with cavities had the strongest cTNT signal followed by organoids with holes, while organoids with intact structures showed little signal (**Figure 1E**). Immunostaining with MYL7 and NRF2F highlighted very few double positive cells (**Figure 1Fi**), suggesting that most of the generated cardiomyocytes (CMs) were indeed ventricular CMs. Simultaneous staining of cTNT and CDH5 revealed a small number of endothelial cells (ECs) lining the inner cell layer of the cavities (**Figure 1Fii**), a similar pattern as endocardial endothelial cells (Endo_ECs) shown in vivo.^27^ Finally, RNA staining of *POSTN*, *WT1*, and *TNNI3* revealed the existence of fibroblasts (FBs) and epicardial cells (EPIs) in the organoids (**Figure 1Fiii, iv**). Notably *POSTN*-expressing cells were observed throughout entire organoids (similar to the FB distribution in vivo), while *WT1* expression was observed only in a small portion of cells, suggesting that EPI or EPI-derived cells only developed in a small region of the organoids.

**Figure 1:**
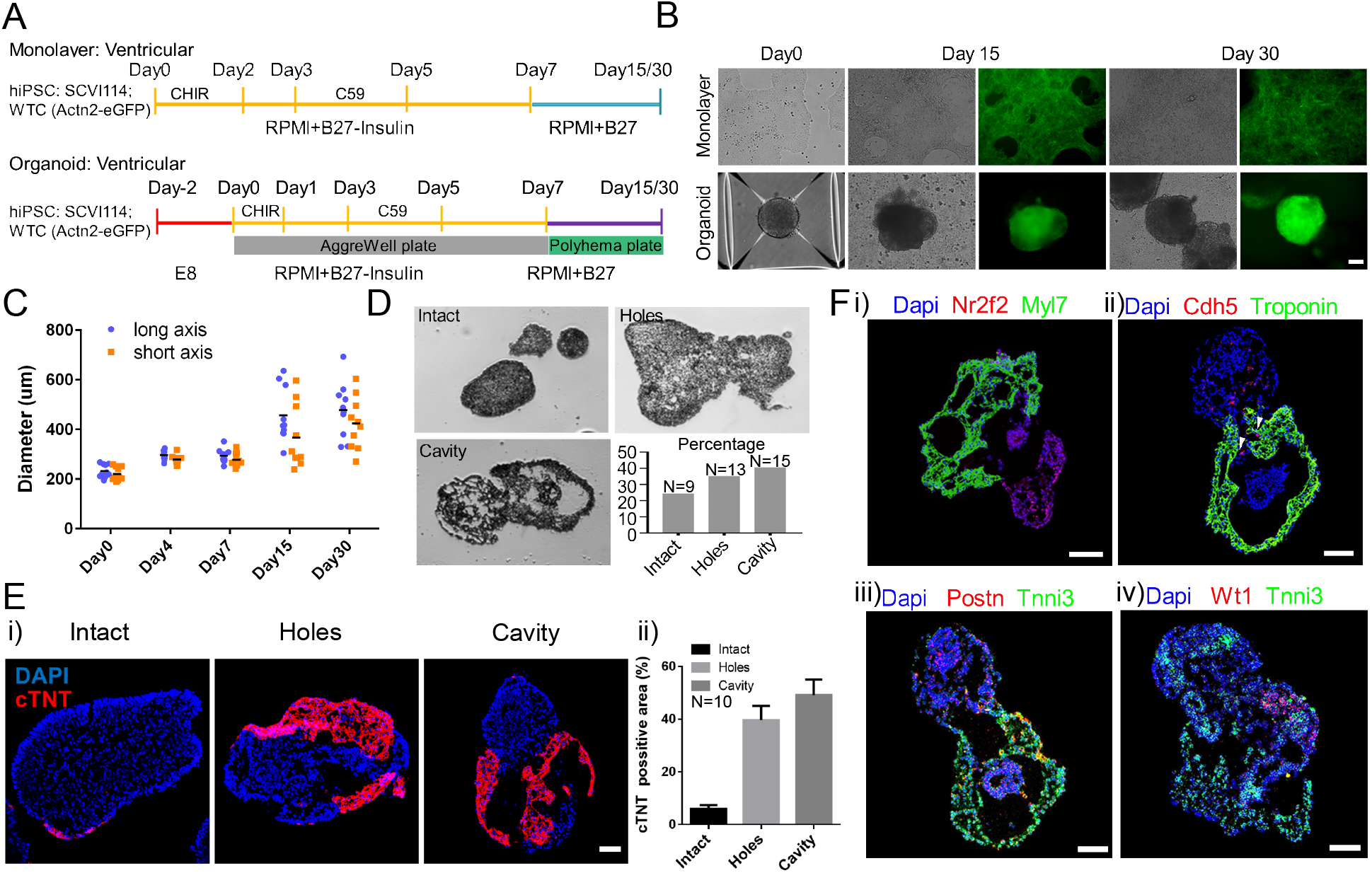
Differentiation and characterization of ventricular heart organoids. (A) Diagram of the ventricular organoid and monolayer cell differentiation workflow. Two cell lines were differentiated in each system. (B) Representative images of the differentiated cells. The GFP signal represents CMs labeled by Actn2-eGFP. (C) Quantification of the organoid diameters from day 0 to day 30. (D) Transverse section analysis revealed three types of organoids. (E) Analysis of the CM areas in the three types of organoids based on cTNT staining (n=10 for each group). (F) In situ expression analysis of cardiac lineage genes with immunofluorescence and RNA in situ hybridizations. Arrowhead points to the Cdh5 positive cells aligning along the inner layer of the cavity. Scale bar=100 um.

Overall, these results showed the heart ventricular organoids we generated captured several important heart developmental characteristics observed in vivo, implying their potential usefulness in studying ventricular cardiogenesis in vitro.

### Transcriptional analysis of ventricular organoids

With the goal of better understanding the cellular and molecular heterogeneity of organoids generated by our protocol, we used single cell RNA sequencing (scRNA-seq) to profile and analyze cells’ transcriptomes. To control for potential batch effects, we employed the MULTI-seq protocol.^23^ In this approach each sample was pre-stained with a unique MULTI-seq sample barcode, and subsequently samples were pooled together and processed with the regular 10X single cell-profiling workflow with minor adaptions.^28^ After Illumina sequencing, data was demultiplexed based on their MULTI-seq barcodes to identify sequencing reads from individual samples (**Figure 2A, S3**).

**Figure 2:**
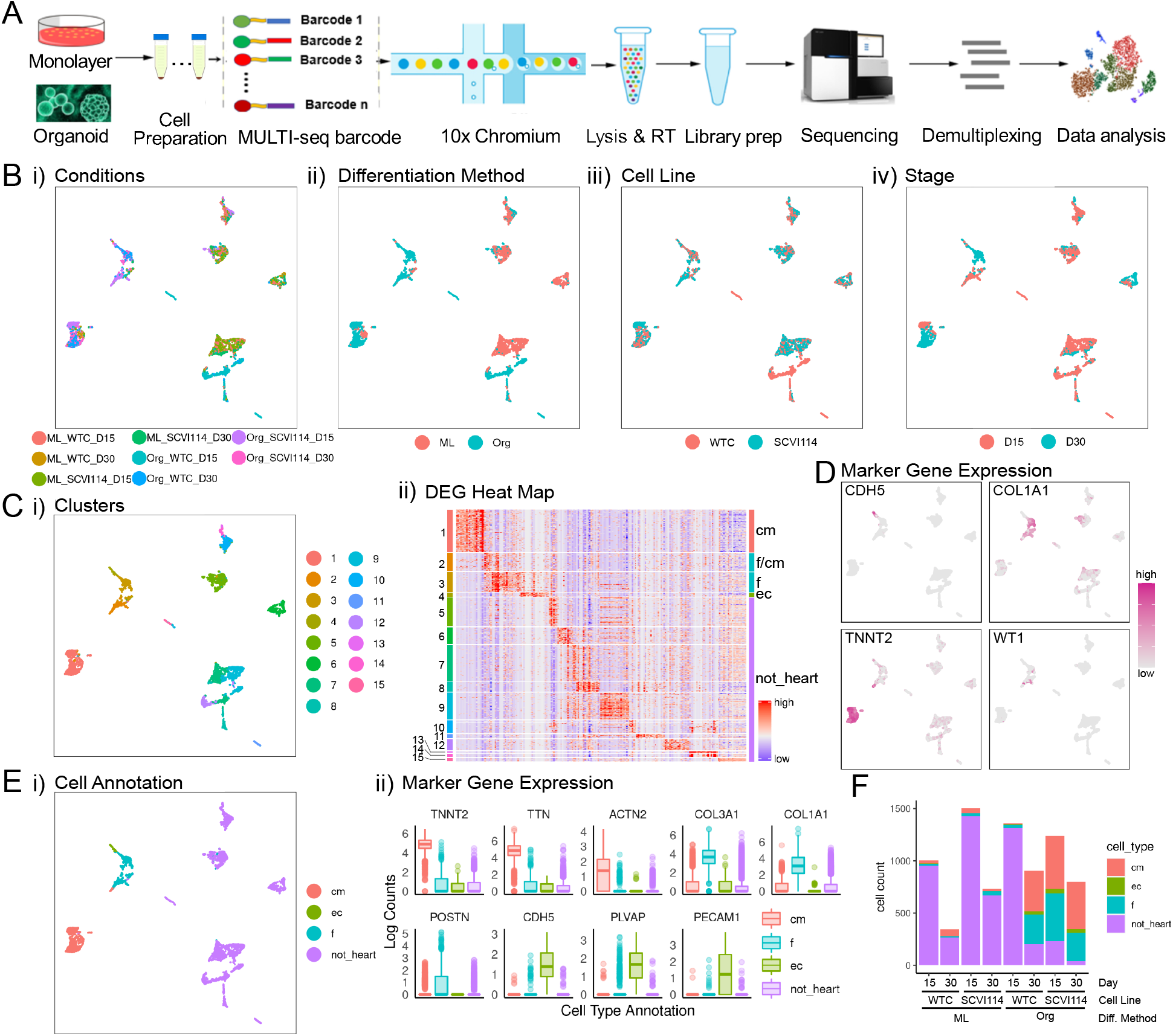
ScRNA-seq analysis of RA-(ventricular protocol) cells. (A) Diagram of the MULTI-seq experimental workflow. (B) UMAP projections of the single cells grouped by i) conditions, ii) differentiation methods, iii) cell lines, and iv) stages. As for all UMAPs in this work, the x-axis is UMAP1 and the y-axis is UMAP2. (C) Unsupervised clustering of the single cells. i) UMAP projection of the clusters and ii) cluster-specific differentially expressed genes with cluster labels (left) and annotated cell types (right). (D) UMAP projections of single cells colored by the expression pattern of representative cardiac lineage genes. (E) i) UMAP projection of single cells grouped by cell type and ii) expression levels of lineage genes in each annotated cell type. (F) The number of profiled cells in each condition colored proportionately by annotated cell type.

Using this approach, we profiled organoid and monolayer differentiated cells (WTC and SCVI114 cell lines) that were generated as described above. After read mapping, demultiplexing and quality control (see **Methods, Figure S2A-B, S3A-B**), we recovered 3,612 cells for the WTC cell line (2,361 at day 15 and 1,251 at day 30) and 4,269 cells for the SCVI114 cell line (2,740 at day 15 and 1,529 at day 30) for further analysis. Unsupervised clustering analysis revealed significant transcriptional differences between cells based on specific combinations of cell line, differentiation protocol (Org vs. ML) and stage (day 15 vs. day 30) (**Figure 2B**). We grouped cells into 15 distinct clusters and identified corresponding unique gene expression signatures (**Figure 2C, Supplemental Table 4**). Together with expression of lineage marker genes (cardiomyocytes: *TNNT2*, *TTN*, *ACTN2*; endothelial cells: *CDH5*, *PECAM1*, *FLVAP*; fibroblasts: *COL1A1*, *POSTN*), we identified three major cardiac cell types (**Figure 2D, 2E**); non-heart-cells did not express cardiac lineage genes but showed expression of genes with roles in brain and kidney development (**Figure 2Eii, Supplemental Table 4**). We found that D15 and D30 organoid cells of the SCVI114 cell line and D30 organoids cells from WTC cell line predominantly differentiated into cardiac cells with only a small percentage specified into non-heart cells; however, monolayer cells and D15 organoid cells from the WTC cell line mostly differentiated into non-heart cells (**Figure 2F**). This observation may be the result of variation in CM differentiation efficiency between experiments. Finally, we also profiled ventricular CMs and non-CMs enriched by FACS based on ACTN2-eGFP expression and identified similar results (**Figure S2C, S3C, S4, S5A-C, Supplemental Table 5**). Overall, scRNA-seq analysis confirmed prior observations about cell function and morphology and showed we were able to generate organoids predominantly consisting of cardiac cell types (CMs, FBs, ECs).

### Generation of atrial heart organoids

In order to generate atrial organoids, we modified our ventricular differentiation work-flow by treating cells with retinoic acid (RA) at cardiac mesoderm and progenitor stages, similar to monolayer atrial differentiation^29^ (**Figure 3A**). Like for ventricular organoids, we differentiated WTC and SCVI114 cell lines and observed beating cells and ACTN-eGFP signal at day 15 and day 30 (**Figure 3B, Video 2**). As before we quantified beating rates and found that atrial organoids showed significantly lower beating rates compared with monolayer cells (**Figure S1B**). Similar to ventricular organoids, we found that atrial organoids grew fast at early stages, then between day 15 and 30 the average size remained similar, but the variance increased markedly (**Figure 3C**). Again, transverse section analysis of the organoids identified three types of internal structures, intact, hole, and cavity; in contrast to ventricular organoids, the hole group is most frequent for atrial organoids (**Figure 3D**). Staining for cardiac troponin (cTNT) revealed that ~66.7% of organoids with cavities, ~23.3% of organoids with holes, and ~15.4% of organoids with intact structures are cTNT positive (**Figure S1E**). Quantification of cTNT positive areas further confirmed that organoids with cavities had the largest CM areas on average (**Figure 3E**). Immunostaining for MYL7 and NR2F2 revealed that many cells stained double positive, suggesting this protocol produced predominantly atrial CMs (**Figure 3F)**. Furthermore, co-staining of CDH5 and cTNT identified a significant proportion of ECs intercalating with CMs, and some lining the cavities (**Figure 3F**). Finally, RNA staining of *POSTN* and *TNNI3* found a large group of fibroblasts; however, most were located separately from CMs (**Figure 3F**).

**Figure 3:**
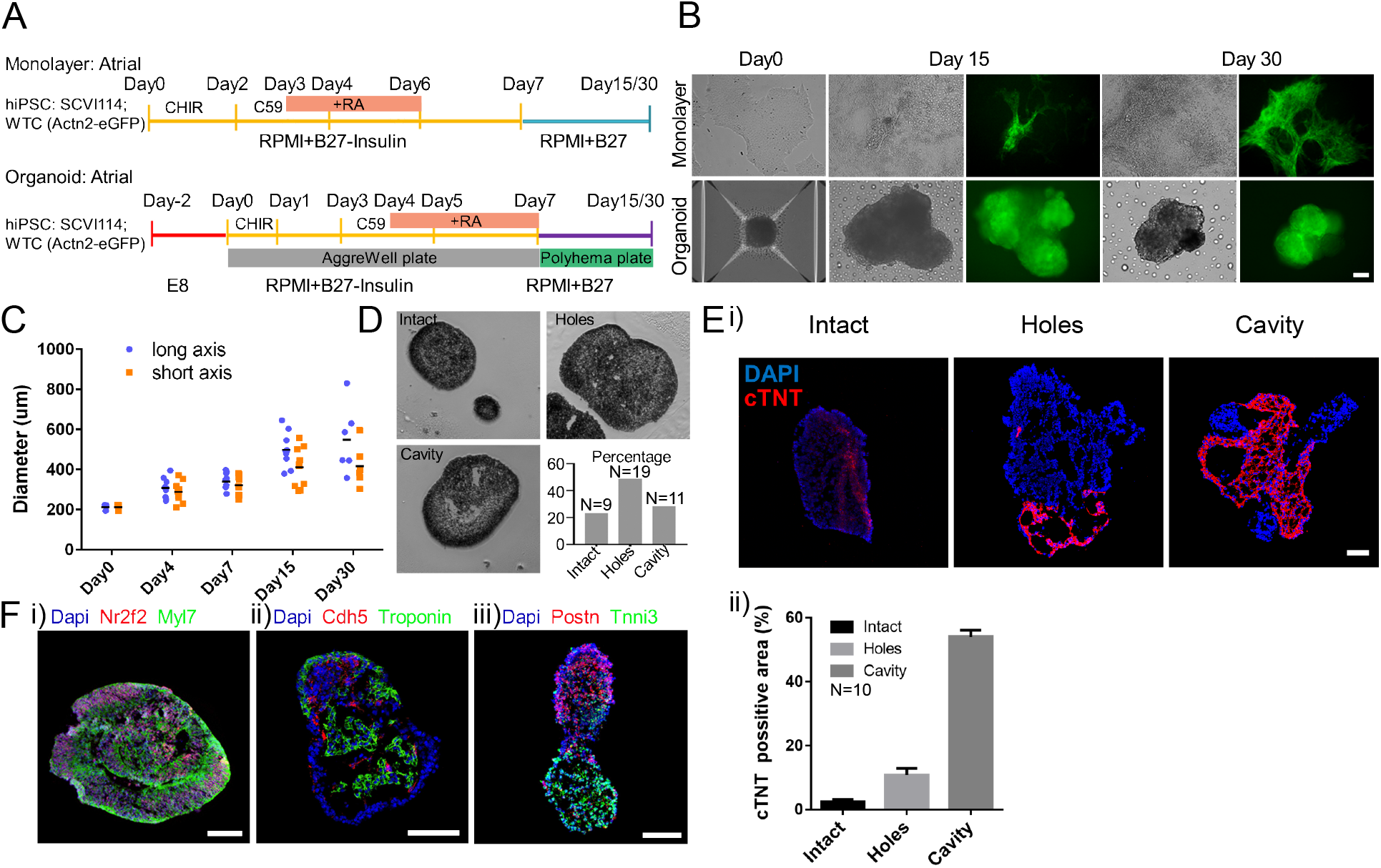
Differentiation and characterization of atrial heart organoids. (A) Diagram of the atrial organoid and monolayer cell differentiation workflow. RA was added to induce the atrial cell lineages. (B) Representative image of the cells at different differentiation stages. (C) The diameter of atrial organoids from day 0 to day 30. (D) Three types of atrial organoids were identified based on their internal structures organoids. (E) Analysis of the CM areas in the three types of atrial organoids based on cTNT staining. (F) In situ expression analysis of cardiac lineage genes in atrial organoids. Scale bar=100 um.

Overall, like for ventricular organoids, we have developed a protocol to generate atrial organoids with the potential to become a valuable in-vitro tool to study atrial cardiac lineage development. Furthermore, together our two differentiation protocols enabled us to compare and contrast atrial and ventricular differentiations in normal and pathological conditions.

### Transcriptional analysis of atrial organoids

Within the same MULTI-seq experiment as ventricular organoids, we also profiled cells from atrial organoid and monolayer differentiations. Differentiated cells at D15 and D30 from WTC and SCVI114 cell lines were analyzed as described above. We recovered 3,551 cells for the WTC cell line (2,329 at day 15, 1,222 at day 30) and 4,042 cells for the SCVI114 cell line (2,060 at day 15 and 1,982 at day 30) for further analysis. Unsupervised clustering followed by projection into two dimensions revealed clear transcriptional differences between differentiation protocols (Org vs. ML) and stages (D15 vs. D30), while differences between the two cell lines (WTC vs. SCVI114) were more subtle and most pronounced in non-heart cells (**Figure 4Ai-iv**). Cells were grouped into 13 clusters and we identified unique expression signatures in each of them (**Figure 4Bi, ii, Supplementary Table 6**). Again, making use of lineage marker genes, we identified CMs, ECs, FBs, and non-heart cells (**Figure 4C, D**). Consistent with what we observed in ventricular organoids, we found that most SCVI114 organoid cells (D15 and D30) and most cells from WTC organoids at D30 differentiated into cardiac cells, whereas WTC and monolayer cells at D15 mainly comprise “non-heart” cells (**Figure 4E**). Finally, we also profiled atrial CMs enriched by FACS based on ACTN2-eGFP expression and found they were a highly pure population of CMs (**Figure S2C, S3C, S4, S5A-C, Supplementary Table 5**). Overall, scRNA-seq analysis confirmed prior observations and showed we were able to generate atrial organoids predominantly consisting of cardiac cell types (CMs, FBs, ECs).

**Figure 4:**
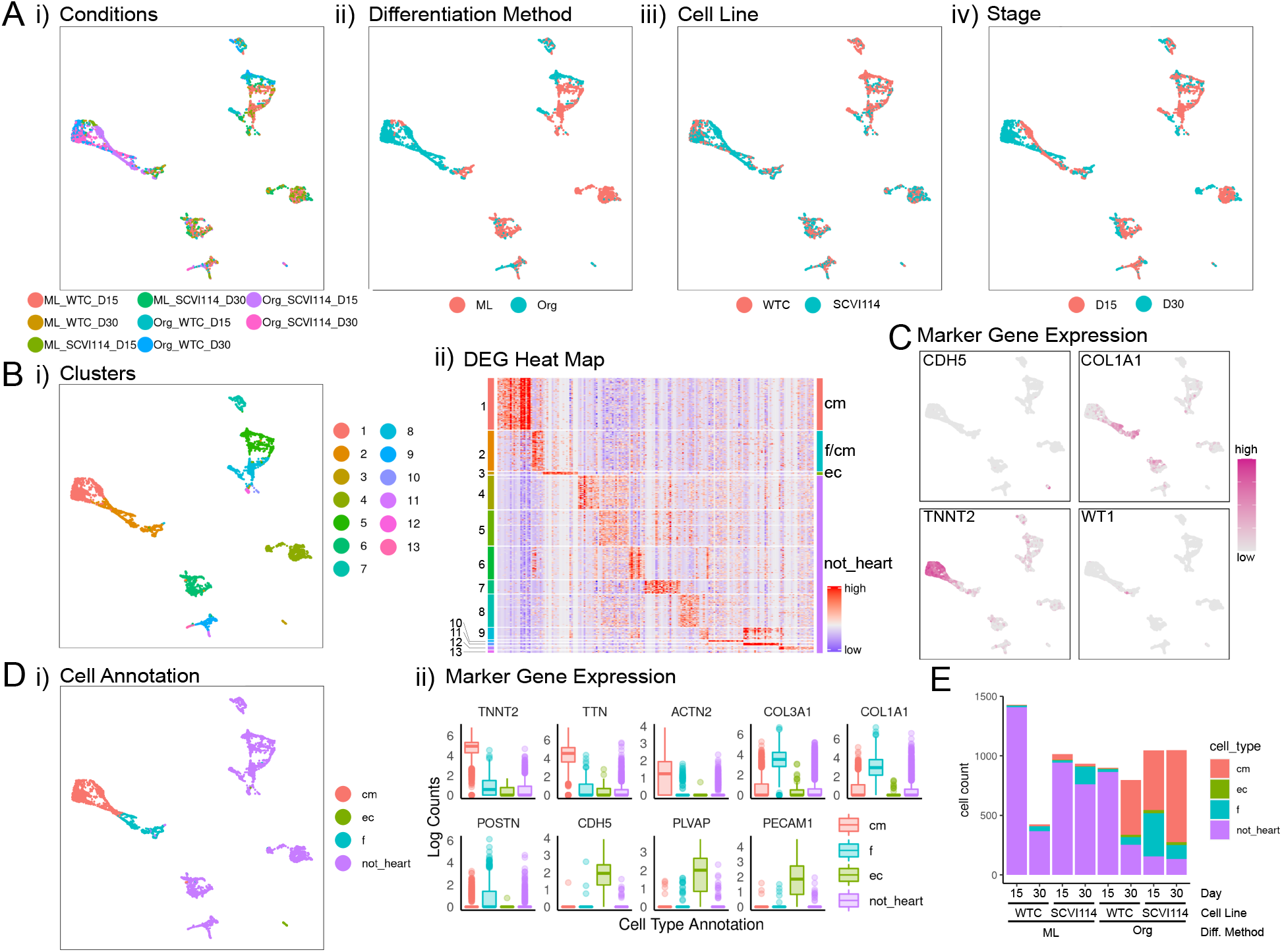
ScRNA-seq analysis of RA+ (atrial protocol) cells. (A) UMAP projections of the single cells grouped by i) conditions, ii) differentiation methods, iii) cell lines, and iv) stages. (B) Unsupervised clustering of the single cells. i) UMAP projection of the clusters and ii) cluster-specific differentially expressed genes with cluster labels (left) and annotated cell types (right). (C) UMAP projections of single cells colored by the expression pattern of representative cardiac lineage genes. (D) i) UMAP projection of single cells grouped by cell type and ii) expression levels of lineage genes in each annotated cell type. (E) The number of profiled cells in each condition colored proportionately by annotated cell type.

### Iterative application of random forests for label transfer from human fetal heart cells

In order to more objectively characterize scRNA-seq data generated from our organoids, we developed a computational approach based on the random forest classification algorithm.^30^ Our goal was to annotate cells from our organoids using published information about (cardiac) cell type, anatomical zone (ventricular vs. atrial) and laterality (left vs. right) from human fetal cells in Cui et al,.^24^ Briefly, to transfer cell type labels we trained a feature selector random forest and a classifier random forest that can then be applied to predict the cell type in test data (our organoid cells, for example). Cells predicted as CMs can then optionally be processed further to annotate anatomical zone and laterality. Finally, our method can perform anomaly detection to filter out cell types that were not present in the training data (**Figure 5A, Figure S6**).

**Figure 5:**
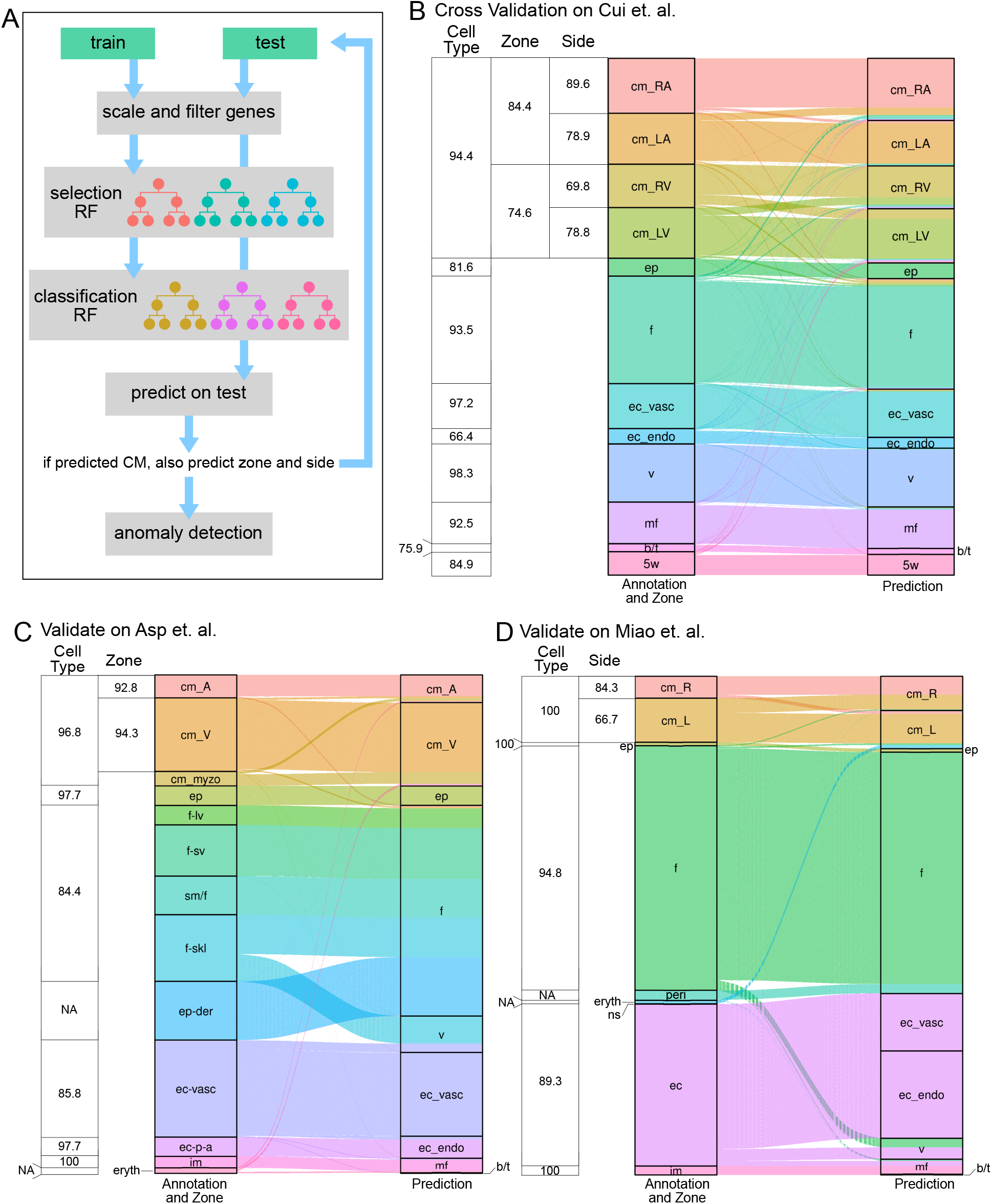
Predicting cardiac cell types, anatomical zones, and laterality. (A) Diagram of hierarchal random forest model. Train and test data are aligned via scaling. Train data is used to derive a random forest model for cell type, which is then applied to test data. For cardiomyocytes the procedure is iterated for predicting anatomical zone and laterality. (B) Sankey diagram of 10-fold cross-validation results on data from Cui et al^24^. Table provides cell type, conditional zone, and conditional laterality/side accuracies. (C) Sankey diagram and accuracies of prediction results for training on the Cui et al. data and prediction on the Asp et al data^25^. (D) Sankey diagram and conditional accuracies of prediction results on Miao et al. data^18^ (training again on the Cui et al. data). Abbreviations: cm = cardiomyocytes, ep = epicardial cell, f = fibroblast, ec_vasc = vascular endothelial cells, ec_endo = endocardial endothelial cells, v = valvar, mf = macrophage, b/t = b/t cell, 5w = undifferentiated cells, cm_myzo = Myoz2-enriched CMs, f-lv = fibroblast-like: (related to larger vascular development), f-sv = fibroblast-like: (related to smaller vascular development), sm/f = Smooth muscle cells / fibroblast-like), f-skl = Fibroblast-like: (related to cardiac skeleton connective tissue), ep-der = Epicardium-derived cells, ec-p-a = Endothelium / pericytes / adventitia, im = immune cells, eryth = erythrocytes, peri = pericyte, ns = nervous system, LA = left atrium, RA = right atrium, LA = left ventricle, RV = right ventricle.

We assessed the performance of this approach in three ways: First, we performed 10-fold cross validation on the Cui et al. data itself. While cross validation guards against overfitting, this is an optimistic scenario because it does not take into account potential differences between training and test datasets. We found that in this setting our approach accurately predicted cell types, anatomical zones and literalities (**Figure 5B**). We noted, though, that performance for cell type prediction worked better (average accuracy = 87.19%) than predicting anatomical zone or laterality (average conditional accuracies of 81.5% and 79.28%, respectively). Second, to take platform differences, variations between laboratories, and other biological variables into account, we used data from Asp et al.^25^, which used the 10X platform (Cui et al. used STRT-seq), to profile atrial and ventricular heart cells (**Figure 5C**). Again, we observed highly accurate prediction of cardiac cell types (average accuracy = 93.73%) and anatomical zones (average conditional accuracy = 93.55%). Interestingly, epicardial derived cells (ep_der), which mostly are fibroblasts, were correctly predicted as such; cardiac skeleton like fibroblasts were predicted as valve cells. Third, we used 10X data from Miao et al.^18^ to assess cell type prediction a second time and, importantly, to assess cross platform laterality prediction. Consistent with our previous results we found cell type prediction highly accurate (average accuracy = 96.82%); prediction of laterality however was only moderately successful, with ~84% of CMs on the right side and ~66% of CMs on the left side correctly classified (**Figure 5D**).

Overall, these results showed that we can use the Cui et al. data set to annotate scRNA-seq data generated on different platforms by different laboratories. We can be highly confident in cell type annotations, confident anatomical zone annotations, and moderately confident in laterality annotations.

### Computational annotation of heart organoid cells’ transcriptomes

Next, we used our computational approach to annotate atrial and ventricular cells at day 30 (**Figure 6**). We found that anomaly detection mainly removed non-heart cells as expected (**Figure S6**), and that cardiac cell types (CMs, ECs and FBs) account for the vast majority of cells and contributed to the major variations in this data set (**Figure 6A**, **B**). Importantly, anomaly detection filtered out few heart cells, which mostly were intermediate cells between fibroblasts and cardiomyocytes (**Figure S6**). Furthermore, remaining non-filtered non-heart cells were predicted as immune cells (macrophages, b/t cells) and fibroblasts. Visual inspection showed that global transcriptional differences between ventricular (RA-) and atrial (RA+) differentiation protocols are most strongly apparent in CMs (**Figure 6Aii, iii**), a trend that was also captured by our anatomical zone predictions (**Figure 6Biii**). Cell type predictions are 91% consistent with our manual cell type annotations, which is also consistent with expectations derived from the validation results as described above. Interestingly, while classification accuracy (in the sense that predictions agree with the differentiation protocols) for atrial (RA+) CMs (82.5%) is within the range of expectation, a fraction of 34.7% of ventricular (RA-) CMs were mis-classified as “atrial”. Further, analyzing CMs in the context of anatomical zone prediction (**Figure 6C**) we found genes that lend support to computational zone predictions (e.g*., MYL7* and *MYH6* (atrial markers)*, MYH7* (ventricular marker)), while others were more consistent with differentiation protocols (*PLN*, *MYH9*, and *MEIS2* (ventricular markers) and *ID3* and *IGFBP5* (atrial markers)). Other marker genes (*NR2F1, NR2F2*) were less clear to interpret. In terms of laterality prediction, we observed that more “left” than “right” CMs, and we found this bias more pronounced for predicted atrial CMs compared with ventricular CMs (**Figure 6E**). For organoids from day 15 we found mostly consistent results (**Figure S8**); cell type predictions were highly accurate (average accuracy = 95.53%), RA+ CMs were largely predicted as atrial (91.2%), however a large fraction of RA- CMs (84.6%) were also predicted as atrial. Also, most CMs at day 15 were not substantially different between RA+ and RA-differentiation protocols (UMAP plot in **Figure S8**), indicating CMs at day 15 may not have matured enough to gain zone identities, which was further supported by the detailed comparative analysis of cells at day 15 and 30 (**Figure S7, Supplementary Table 12**).

**Figure 6:**
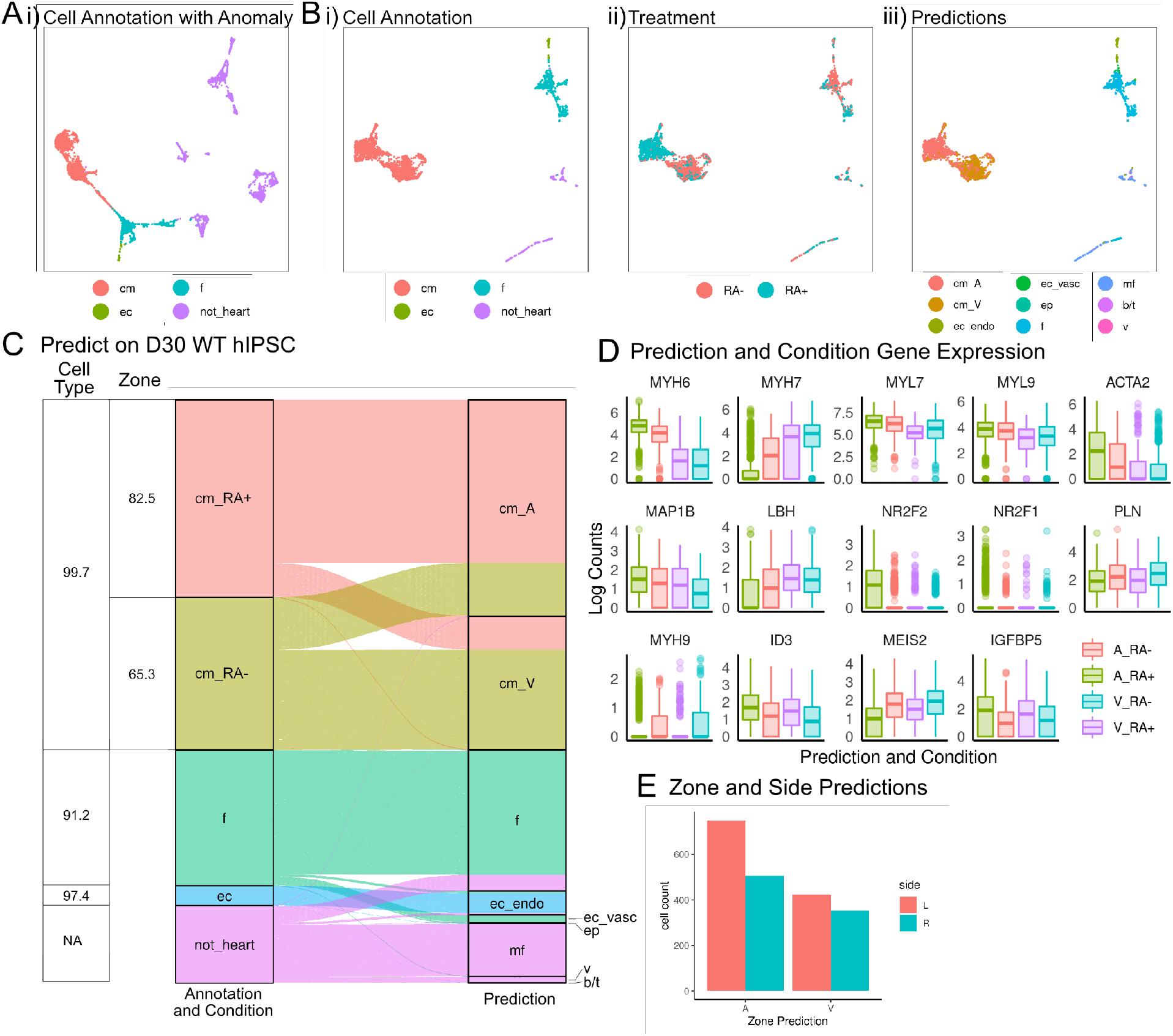
Prediction of wild type RA- and RA+ cells using the validated hierarchal random forest model. (A) UMAP projections of RA- and RA+ single cells at day 30 grouped by cell type annotation. (B) UMAP projections of wild type RA- and RA+ single cells at day 30 with Anomalies removed grouped by i) cell types, ii) treatment, and iii) predicted cell types and zone. (C) Sankey diagram of prediction results of wild type RA- and RA+ cells at day 30 (Anomalies removed). Table provides cell type and conditional zone accuracies. (C) Expression levels of genes used to make prediction decisions that either correlate with the predictions (A and V) or experimental conditions (RA- and RA+). (D) Bar plot of side predictions in WT cells at day 30.

We also analyzed the cardiac cells enriched by FACS. We again found cell type predictions highly accurate (average accuracy = 94.4%), as were anatomical zone predictions: 94.7% of RA+ CMs (atrial protocol) were predicted as atrial CMs, and 62.9% of RACMs (ventricular protocol) were predicted as ventricular CMs (**Figure S5, Supplementary Table 5**).

Overall, our automatic predictions achieved high accuracy in cell type annotations and highlighted that zone identities were more established at day 30 compared to day 15 in our organoid differentiation systems. These results enabled us to use this computational phenotyping approach to compare wild type and genetically modified organoids.

### Generation of hiPSC lines and organoids carrying a genetic variant associated with Ebstein’s Anomaly

The homeobox-containing transcription factor *NKX2-5* plays critical roles in embryonic heart development.^12,31^ Notably, *NKX2-5* knockout mice die at E10.5 with only two heart chambers, both with atrial identities as reported by ATLAS-seq predictions.^17^ Furthermore, a single nucleotide variant in the *NKX*2-5 gene locus at the 673^th^ nucleotide converting the 188^th^ amino acid from Aspartate (N) to Lysine (K) was associated with Ebstein’s Anomaly, a congenital heart defect diagnosed with atrialized right ventricle and abnormal tricuspid valve.^11,32^ We next used our two differentiation protocols for producing atrial and ventricular organoids, together with CRISPR/Cas9 technology, to characterize and study the effects of the above-mentioned variant.

In order to do so, we produced an isogenic line introducing this mutation into the WTC line using a single-stranded oligodeoxynucleotide (ssODN) based CRISPR/Cas9 strategy and selected two clones (PM28 and PM52) for differentiation. As a control we created a line where the first exon of *NKX2-5* was deleted (Del33) using a pair of sgRNAs (**Figure 7A**). After differentiation we observed ACTN2-GFP signal and beating cells in ventricular and atrial organoids at day 30 (**Figure 7B**). Additionally, in the large deletion line we observed significant reduction of Nkx2-5 immunostaining signal in the differentiated organoids (**Figure 7C**). We note that, compared with wild type organoids, we also observed significant reduction of beating rates in the mutant atrial organoids, but not in ventricular organoids (**Figure S9**).

**Figure 7:**
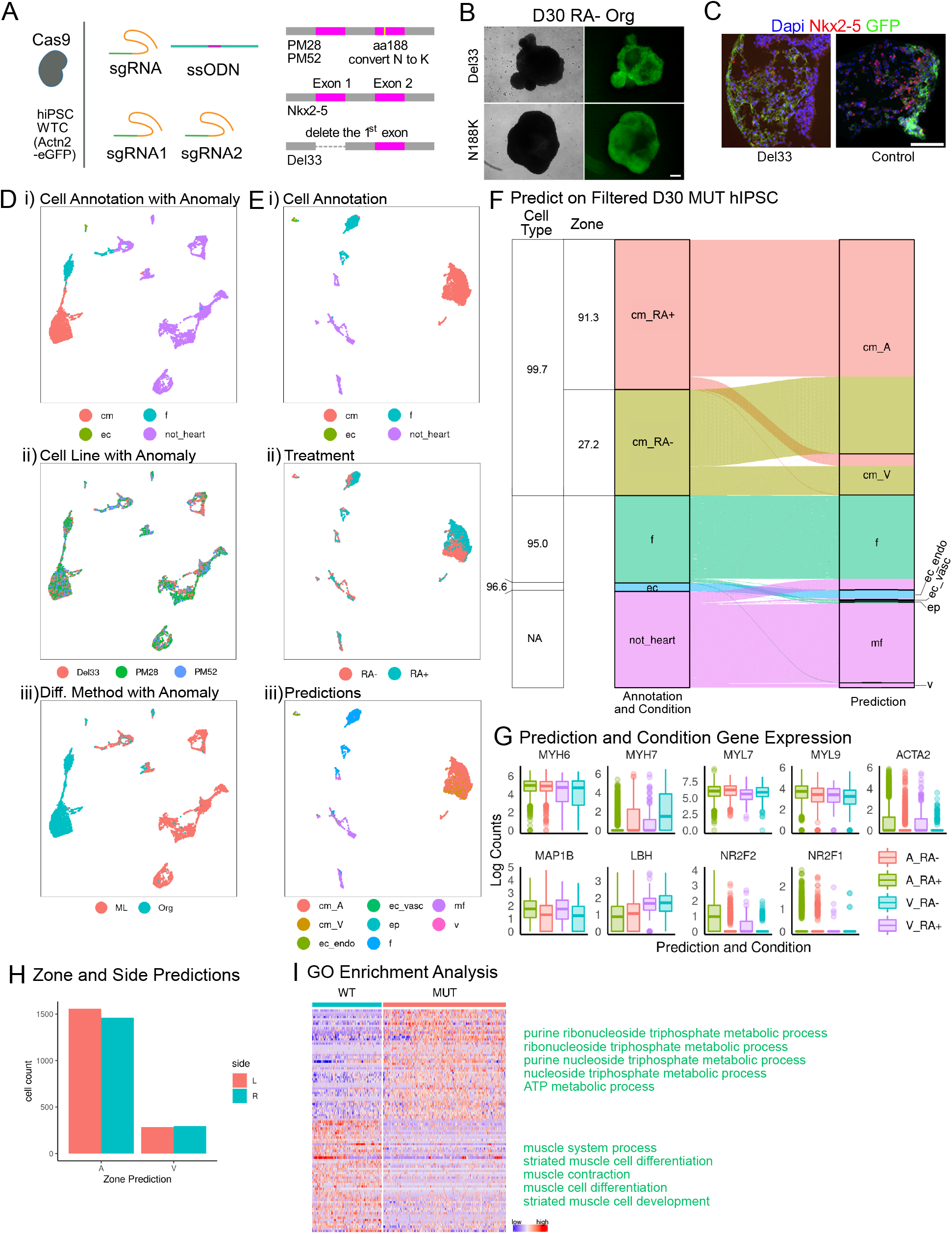
Study of EA defects in a mutant isogenic line using random forest predictions. (A) Diagram of the strategy to generate an isogenic line carrying an EA associated mutation and a control line with a large deletion on Nkx2-5 (Del33). (B) Representative organoids differentiated from the two mutants at day 30. (C) Nkx2-5 expression significantly reduced in the Del33 line derived organoids. (D) UMAP projections of mutant RA- and RA+ single cells at day 30 grouped by i) cell types, ii) cell line, and iii) differentiation method. (E) UMAP projections of day 30 mutant RA- and RA+ single cells with Anomalies removed grouped by i) cell types, ii) treatment, and iii) predicted cell types and zone. (F) Sankey diagram of prediction results of mutant RA- and RA+ cells at day 30 (Anomalies removed). Table provides cell type and conditional zone accuracies. (G) Expression levels of genes used to make prediction decisions that either correlate with the predictions. (H) The results of side predictions in mutant cells at day 30. (I) Gene pathways that differentially enrich in the wild type and mutant (predicted) CMs. Scale bar=100um.

### Transcriptional analysis of mutant heart organoids

Within the same MULTI-seq setup as discussed above we sequenced organoids from the PM28, PM52 and Del33 cell lines at day 30. After bioinformatics processing and quality control, we recovered 5,596 (PM28), 3,465 (PM52) and 4,367 (Del33) cells for down-stream analysis. Unsupervised clustering and (manual) lineage gene analysis revealed major cardiac cell types (**Figure 7Di, 7Ei, S10**). Next, we used our cell type classification approach to automatically annotate cell types in this dataset. Consistent with previous results we found that anomaly detection predominantly filtered out non-heart cells, and that predicted cell type annotations were highly consistent with our manually inferred cell types (average accuracy =95%, **Figure 7D, E, F**). We did not observe pronounced differences between the three mutant lines (PM28, PM52, Del33, **Figure 7Dii**), but found cells from organoids were largely different from monolayer-differentiated cells (**Figure 7Diii**), a signal that is driven by most monolayer-differentiated cells likely not being heart cells (**Figure 7Di, iii**). While, like wild type cells, global transcriptional differences between ventricular (RA-) and atrial (RA+) differentiation protocols manifest mostly in CMs (**Figure 7Ei, ii**), the strong distinction between RA- and RA+ treated cells that was observed in wild type CMs was lost. We also found a significant fraction of RA-(i.e., ventricular) CMs annotated as “atrial” by our label transfer procedure. While we did observe this type of “misclassification” in the wild type cell lines, the “misclassified” fraction of cells in mutants significantly increased (72.8% here vs. 37.4% for wild type organoid cells at day 30, **Figure 7F** and **Figure 6B**). Similar to before we found atrial genes (including *MYL7*, *MYH6, MYL9*) highly expressed in predicted atrial CMs from organoids differentiated with the ventricular protocol (RA-) (**Figure 7G**). Like in the wild type cells, we observed laterality bias in CMs (**Figure 7H**); however, for the modified lines we observed slightly more “right” ventricular cells, which was the opposite of wild type cells (**Figure 6D**).

Finally, when comparing gene expression between wild type and mutant hiPSC-derived cells, we found that there were significantly more differentially expressed genes (DEGs, fdr<0.001) between wild type and mutant cells in CMs (166), compared with FBs (21), and Endo_ECs (2) (**Supplementary Table 7**). In order to rule out the possibility that differences in cell type abundances account for this observation (we find 279 Endo_ECs, 2,158 FBs, 5,628 CMs), we down-sampled CMs to either 279 cells (like Endo_ECs) or 2,158 cells (like FBs) 200 times and re-calculated DEGs. This procedure confirmed more DEGs in CMs compared to other cell types (**Figure S12**). This shows that the mutation of *NKX2-5* mainly affects CMs and we see little evidence for effects on Endo_ECs and FBs. Focusing on CMs, Gene Ontology enrichment analysis of DEGs, highlights striated muscle development-related biological processes amongst the top down-regulated genes (low expression in mutant CMs), while the most significantly enriched terms in up-regulated genes (high expression in mutant CMS) highlight processes related to metabolism and energy (**Figure 7I, Figure S11**).

Overall, these results demonstrate that we successfully generated ventricular and atrial organoids with a specific genetic variant associated with Ebstein’s Anomaly. Our classifier predicted most mutant CMs differentiated in the ventricular (RA-) protocol as atrial CMs, recapitulating the atrialized ventricular defects in EA. Furthermore, our observations suggested that the *NKX2-*5 mutation predominantly impacts cardiomyocytes, with genes related to striated muscle differentiation showing weaker expression in mutant CMs, while genes related to metabolism and energy production appear up-regulated. This for the first time provides clear evidence that the EA-associated variant (c.673C>A)^11^ affects the expression of energy-related and key heart muscle genes during (in vitro) cardiogenesis, building confidence in this particular variant’s relevance and laying the foundation for more detailed disease models of EA.

## Discussion

In this study, we generated ventricular and atrial heart organoids and used scRNAseq in combination with MULTI-seq sample pooling to obtain transcriptional profiles at single cell resolution.^23,28^ We established a machine learning label transfer method that allowed us to leverage annotations (cell type, anatomical compartment, laterality) from primary human fetal cells, and we used this approach to characterize cells differentiated with our organoid systems. Finally, we used this experimental and computational combination to compare differentiated organoids from wild type cell lines with organoids carrying a genetic variant associated with EA, effectively establishing an in vitro hiPSC model for this type of congenital heart defect.

We find that our organoid systems recapitulated the microenvironment of human developing hearts by self-assembling into chamber-like structures. We note that this type of three-dimensional approach has advantages when studying heart developmental processes, especially chamber formation, compared to monolayer and co-differentiated microtissue systems.^33,34^ We note that along the same lines Hofbauer et al^3^. recently reported that the addition of VEGF to cardioids (a similar type of cardiac organoid) can lead to the development of endothelial cells that comprise the entire inner layer of chambers, essentially equivalent to the in vivo anatomical pattern observed in endocardial endothelial cells.^3^ With this study and our work generating specific cell types in environments resembling their in vivo anatomical compartments, it will be interesting to establish differentiation protocols in the future that mimic the in vivo localization of other cardiac cell types, like epicardial cells and vascular endothelial cells.

Based on the sarcomere lengths and scRNA-seq data, we found the hiPSC derived CMs in both organoid and monolayer systems were relatively immature compared to primary cells. Additionally, we found the CMs in organoids were slightly less mature compared with CMs in monolayer system (**Figure S7, Supplementary Table 12)**. However, since it has been reported that co-culture of CMs and other cardiac cells (FBs, ECs) can significantly improve CM maturation, we believe that organoid cells may further mature after long-term culture.^35^ Furthermore, cells often become more mature after extensive culture, as has been reported for other types of organoids such as brain.^36^ We note, though, that this phenomenon is likely tissue- or organ-specific, because kidney organoids did not mature after further culturing, but instead showed higher cell death rate.^37^ However, it is known that in vitro differentiated CMs can mature significantly after being transplanted into live organisms.^38^ While it would be challenging to transplant generated heart organoids to replace the heart of model organisms, it may be feasible to transplant them into other parts of animal models, like mice or zebrafish, to study their maturation.

While in silico phenotyping of organoid cells performed well overall, the experimental setup with different platforms for training (STRT-seq) and testing/application (10X) is clearly not optimal, and generating more similar test/training data in future experiments will likely increase accuracy and reliability of computational phenotyping. In addition, making use of spatial transcriptomics approaches to increase resolution and confidence of annotations with regard to anatomical zone and laterality, without the need for tissue dissections, would yield an increased chance of capturing more subtle transcriptional spatial features. When predicting anatomical zones for wild type organoid cells, the vast majority of predictions agreed with the differentiation protocol (atrial = RA+, ventricular = RA-, see **Figure 6C**). However, for a smaller group of cells the protocols and predictions mismatch, that is RA- differentiated cells were predicted “atrial” and RA+ differentiated cells were “ventricular”. Further investigating those mismatching cells yielded zone-specific genes that support zone predictions (e.g., atrium-specific genes *MYL7* and *MYH6* are high in RA- differentiated but “atrial” predicted cells) and others more in-line with the differentiation protocols (e.g., ventricle-specific genes *PLN*, *MEIS2*). In the future it will be interesting to further investigate these genes, and more specifically elucidate their relation to RA, the only difference between the atrial and ventricular differentiation protocols. However, to assess whether direct regulation by RA plays a role will require further experiments, like applying ChIP-seq or derivative technologies in generated organoids.^39^

We also noted that the fraction of heart cells we recovered by scRNA-seq varied between the monolayer (low fraction of heart cells recovered) and organoid protocols (high fraction of heart cells recovered), see **Figures 2F**,**4E**. Exception to that rule were the WTC organoids at day 15. Since monolayer protocols have been reported to generate cardiomyocytes with high efficiency, we believe this observation may be specific to this batch rather than being representative of the monolayer approach. Therefore, we interpret the data in the sense that it shows our organoid protocol to be efficient.

Our protocols also allowed us to produce genetically modified cell lines carrying a mutation associated with EA, and compare resulting organoids with wild type differentiations. We found that genes down-regulated in mutant organoids were associated with striated muscle differentiation, while mutant up-regulated genes were often related to energy and metabolism, which provided the first (in vitro) characterization of molecular effects of the c.673C>A mutation and may constitute a first step towards more detailed models of the contribution of this genetic lesion towards EA. We note, however, that in addition to atrial/ventricular CM lineage defects, EA patients also present with tricuspid valve defects. Our current system does not model this aspect, but we can extend our approach (differentiation system and cell type / anatomical zone computational modeling) to include and focus on valve-related cells in the future. Furthermore, EA is known to be genetically multigenic,^9,40^ and our general approach of in vitro modeling together with computational phenotyping can be applied to other EA-associated genetic variants to gain systematic insight into the disease.

In summary, in this work we have established chamber-specific differentiation protocols for heart organoids, and we showed that in combination with scRNA-seq profiling of organoid cells this system is a useful model for investigating genetic lesions at the *NKX2-5* locus associated with EA. While it was necessary to focus on zone/chamber specificity (atrial vs. ventricular) in this context, our approach can be repurposed to focus on laterality (left vs. right), which would be interesting in the context of CHDs with known laterality phenotypes, such as heterotaxy and hypoplastic left or right heart syndrome.

## Materials and methods

### Maintenance of hiPSC lines

WTC line with ACTN2-eGFP transgene (Coriell catalog: GM25256) and SCVI114 line (Gift from Stanford CVI) were maintained in completely defined albumin-free E8 medium (DMEM/F12 with L-glutamine and HEPES, 64μg/ml L-Ascorbic Acid-2-phosphate, 20μg/ml insulin, 5μg/ml transferrin, 14ng/ml sodium selenite, 100ng/ml FGF2, 2ng/ml TGFb1) ^41^ on Matrigel (Corning, CB40230A) coated tissue culture plates. Medium was changed daily and routinely passaged every three to four days using 0.5 mM EDTA solution (Invitrogen, 15575020). 10 μM ROCK inhibitor Y27632 (Selleckchem, S10492MG) was supplemented to the medium during cell passaging.

### Monolayer cardiac differentiation

Monolayer cardiac differentiation was carried out following a published protocol^42^. Briefly, RPMI 1640 media (Corning, 10040CVR) was used as the basal medium in the entire differentiation process. B-27 Supplement minus Insulin (Gibco, A1895601) was supplemented to the medium for the first 6 days and B-27 Supplement (Gibco, 17504044) was used afterwards. The small molecule inhibitor of GSK3β signaling, CHIR99021 (Selleckchem, S292425MG) was used in the first two days of differentiation and Wnt signaling inhibitor C59 (Selleckchem, S70375MG) was added from day 3 to day 4. To differentiate atrial cells, 1 uM retinoic acid (Sigma-Aldrich, R2625) was added from day 3 to day 6 as described Previously ^29^.

### Cardiac organoid differentiation

The cardiac organoid differentiation procedure was adapted from a protocol described Previously ^2^. Briefly, 1.5×10^6^ hiPSCs were seeded in each well of AggreWell™800 plates (STEMCELL, 34815) according to the manufacturer’s instructions. The cells were assembled into 3D structure by culturing in E8 medium for 2 days (day -2 to day 0). From day 0 to day 6, cells were cultured in RPMI supplemented with B-27 minus insulin. CHIR99021 at a final concentration of 11μM was used at day 0 and lasted for 1 day. From day 3 to day 5, cells were treated with C59 at a final concentration of 5 μM. The cell aggregates were transferred to 5% Poly(2-hydroxyethyl methacrylate) (Sigma-Aldrich, P3932) coated tissue culture plates at day 7 and cultured in RPMI with B27 supplement until the end of differentiation. Fresh medium was changed every 3 days until tissue harvest. To differentiate organoids, 1 uM retinoic acid was added from day 4 to day 7.

### Generating Nkx2-5 mutant hiPSC lines

To generate Nkx2-5 loss of function mutants, a pair of single-guide RNAs (sgRNAs) (**supplemental Table 1**) were used to target the first exon of Nkx2-5 gene. The sgRNAs were cloned into pSpCas9(BB)-2A–GFP (PX458) vector and transfected into WTC hiPSCs with nucleofector. Specifically, about 8 × 10^5^ iPSCs were transfected with 5 μg of plasmids with Lonza Human Stem Cell Nucleofector Kit 1 (Lonza, VPH-5012) on a Nucleofector 2b device (Lonza, AAB1001). After FACS sorting and PCR genotyping of multiple iPSC clones, we identified the clones with the deletion of first Nkx2-5 exon and further expanded them for cardiac differentiations.

Besides, we introduced a single nucleotide mutation into the Nkx2-5 gene locus using a ssODN based CRISPR strategy. We co-transfected a ssODN and sgRNA (**supplemental Table 1**) to convert the 673^th^ nucleotide from C to A, which led the protein change at 188 amino acid from Asn to Lysin. After clone picking and genotyping the clones by sanger sequencing, we identified the positive clones and further expanded them for differentiation.

### Single cell isolation

Cardiac cells from monolayer differentiation culture at day 15 and day 30 were washed twice with PBS and incubated with TrypLE Express (Life Technologies, A1217702) for 15 min at 37 °C. Cells were collected by centrifuge at 300 g for 5 min and washed once with HBSS−/− (Ca^2+^/Mg^2+^ free). The cells were further resuspended in 1ml B27 and filtered through a 40 μm filter (Corning, 431750). After that, the cells will be ready for FACS sorting or directed used for scRNA-seq.

The cardiac organoids were collected and washed twice with HBSS−/− (Ca^2+^/Mg^2+^ free) before being incubated with 0.25% Trypsin/EDTA at 37 °C for 5 min. After that, a collagenase HBSS+/+ mixture with 10 mg/ml of collagenase A (Sigma-Aldrich, 10103586001), 10 mg/ml of collagenase B (Sigma-Aldrich, 11088815001), and 40% FBS (Gibco, 26140079) was added to the digestion solution and gently pipetted until the organoids were completely dissociated. The cells were then spun down at 300 × g for 5 min, resuspended in 1ml of RPMI/B27 medium, and filtered through a 40 μm filter.

### MULTI-seq barcoding

First batch (FACS sorted). The cells from monolayer differentiations and organoids were washed once with FACS buffer (HBSS−/−,10% FBS) and resuspended in 1 mL of FACS buffer with 10 μM ROCK inhibitor. After FACS sorting based on GFP expression (BD, FACSAria™ III), both GFP positive and negative cells were collected separately for scRNA-seq. Each sample was stained with a unique MULTI-seq barcode following a published protocol with minor modifications^23^. Briefly, the cells were washed twice with PBS (Ca^2+^/Mg^2+^ free) and resuspended in 180 μl PBS (Ca^2+^/Mg^2+^ free). 20ul of 2 μM sample specific Anchor/Barcode (**Supplemental Table 2**) was then added to each sample and incubated for 5 min on ice. After that, 20 μl of 2 μM Co-Anchor solution (**Supplemental Table 2**) was added and kept on ice for another 8 mins. The samples were then washed once and resuspended in ice cold PBS with 1% BSA. The cell numbers were counted before pooling the samples together.

Second batch (Not sorted). All cells including the monolayer differentiated cells and organoid differentiated cells (WTC and SCVI111 line differentiated in atrial and ventricular differentiation protocols at differentiation day 15 and 30, and Nkx2-5 mutant line in atria and ventricular differentiation protocols at day 30) were prepared as single cells. The cells in each sample were stained with MULTI-seq barcode following the same procedure as the first batch of cells. Afterwards, the cells were counted and pooled together for scRNAseq.

### Library preparation and single-cell mRNA sequencing

The pooled cells were captured in 10X Chromium (10X Genomics, 120223) by following the single cell 3’ reagent kits v3 user guide. Briefly, cells were loaded into each chip well to be partitioned into gel beads in emulsion (GEMs) in the Chromium controller. We targeted for 25,000 cells in each chip well and profiled one well for the first batch experiment and two chip wells for the second experiment. The cells were then lysed and barcoded reverse transcribed in the GEMs. After breaking the GEMs and further cleanup and amplification, the cDNA was enzymatically fragmented and 3’ end fragments were selected for library preparation. After further processing including end repair, A-tailing, adaptor ligation, and PCR amplification, a string of sequences including sample index, UMI sequences, barcode sequences, and sequencing primer P5 and P7 were added to cDNA on both ends. The libraries were sequenced on Illumina HiSeq × platform.

### Bioinformatics analysis

#### Data processing and quality control

Alignment and quantification of UMI counts for endogenous genes were performed using the cellranger count pipeline of the Cell Ranger software (version 3.1.0). We used the human reference genome (GRCh38.p12) and arguments --chemistry= SC3Pv3 and --expect-cells as 10,000 or 25,000, depending on the specific library. For sample demultiplexing, we used the R package deMULTIplex (version 1.0.2, https://github.com/chris-mcginnis-ucsf/MULTI-seq) which consists of alignment of the MULTI-seq sample barcode read sequences to the reference MULTI-seq sample barcodes followed by sample classification into doublets and singlets. Multiple quality control (QC) metrics were calculated using the R package scater (http://www.bioconductor.org/packages/release/bioc/html/scater.html), and cells with total library size >=2000, number of detected genes >=1000 and <= 8000, and <=30% percentage of mitochondrial reads were considered. To account for doublets with the same MULTI-seq barcode we used the scds R package (https://github.com/kostkalab/scds) as described below. We focused on genes with one or more count in at least five cells (assessed for each batch separately) and calculated log-normalized counts using the deconvolution method of the scran R package (https://bioconductor.org/packages/release/bioc/html/scran.html). There are two batches in the unsorted data, so multiBatchNorm from the package batchler (https://bioconductor.org/packages/devel/bioc/html/batchelor.html) was used to perform scaling normalization so that the size factors are comparable across batches. Next, clustering, dimensionality reduction, and cell type annotation was performed separately on wildtype atrial, wildtype ventricular, and mutant groups. The top 2000 highly variable genes were identified using the modelGeneVar function (scran R package). Using these genes, 50 principal components were calculated (runPCA, scater R package) and used to generate UMAP plots (runUMAP, scater R package) and to build a shared nearest neighbor graph followed by walktrap clustering (cluster_walktrap, igraph R package, https://github.com/igraph/igraph) as outlined by Amezquita et al.^43^

For FACs sorted data no batch correction was needed, so clustering was performed by building a shared nearest-neighbor graph using the first 25 first principal components for each cell; we used Jaccard weights and the Louvain clustering algorithm from the igraph package with steps = 10 parameter. The R package ComplexHeatmap (http://www.bio-conductor.org/packages/release/bioc/html/ComplexHeatmap.html) was used to generate gene expression heatmaps and findMarkers (scran R package) with the fdr = .001 parameter was used to get inter-cluster differentially expressed genes. Finally, cell type annotations were manually resolved using cluster expression patterns of the following genes: *TNNT2, ACTN2, TNNI3, TTN, MYH6, NR2F2, MYL2, MYH7, COL1A1, DCN, SOX9, POSTN, WT1, TBX18, ALDH1A2, LRRN4, CSF1R, TPSAB1, CD3D, GIMAP4, PECAM1, CDH5, TIE1, NPR3, PLVAP, FOXC1, FABP4, CLDN5, HEMGN, HBA-A1, HBA-A2, C1QA*. In the rare case where a cluster expresses marker genes for more that one cell type, iterative clustering was performed to resolve cell types.

#### Computational annotation of multiplets in a MULTI-seq workflow

The MULTI-seq approach identifies multiplets based on occurrence of more than one MULTI-seq cell barcode. By design, this approach cannot identify mutiplets comprised of cells with identical MULTI-seq sample barcodes. We use computational multiplet identification (scds) to identify this type of *“within-sample”* multiplet computationally. Broadly, we use MULTI-seq data to estimate the fractions of within-sample and between-sample multiplets and use them to determine the number of within-sample multiplets that we annotate computationally.

Specifically, in our approach we assume the overall fraction of cells^1^ being multiplets, *p*_*m*_, is comprised of within-sample multiplets (*p*_*w*_, with the same MULTI-seq barcode) and between-sample multiplets (*p*_*b*_, with distinct barcodes) and no other contributions: *p*_*m*_ = *p*_*w*_ + *p*_*b*_ = *p*_*m*_(*p*_*w*_/*p*_*m*_ + *p*_*b*_/*p*_*m*_) = *p*_*m*_π_*w*_ + *p*_*m*_π_*b*_, where π_*w*_ and π_*b*_ denote the fraction of multiplets being within-sample and between-sample, respectively; also: π_*w*_ + π_*b*_ = 1. We then use the following ansatz, where the fraction of different types of multiplets is proportional to the abundance of constituent cells (we only focus on doublets and assume higher-order multiplets to be rare): 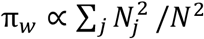 and π_*b*_ ∝ ∑_*(i,j)*,*i*>*j*_ *N*_*i*_*N*_*j*_/*N*^2^ where *N* denotes the overall number of cells and *N*_*k*_ the number of cells with MULTI-seq/sample barcode *k*. Utilizing estimates of these quantities obtained by demultiplexing MULTI-seq data and the constraint that π_*w*_ and π_*b*_ sum to one we obtain estimates 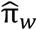 and 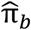.

Let *D*_*m*_ be the number of MULTI-seq annotated between-sample multiplets. We have *D*_*m*_ = *Np*_*m*_π_*b*_ and therefore *p*_*m*_ = *D*_*b*_/(*N*π_*b*_) and plugging in 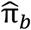 yields 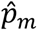, an estimate for the fraction of doublets in our data set; note that 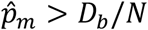, the fraction of multiplets obtained from the MULTI-seq data alone. The “missing” number of within-sample multiplets is then estimated as 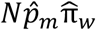, determining the number of doublets we annotate using scds in addition to the between-sample doublets annotated by MULTI-seq.

#### Random forest based cell type classification across data sets

##### Training data

We used the single cell RNA-seq data of Cui et al.^24^ with cell type, anatomical zone, and laterality annotations in order to train a random forest classifier that generalizes to other data sets. Data and cell annotations were downloaded from GEO (GSE106118)^24^. Iterative clustering was used to further resolve the annotated ECs into Endo_ECs and vascular ECs based on the expression of Endo_EC markers (*NPR3*, *PLVAP*, *FOXC1*) and Vasc_EC markers (*FABP4* and *CLDN5*). Additionally, iterative clustering was used to reannotate a subcluster of “5w” cells as epicardial cells based on the expression of *WT1*, *TBX18*, *ALDH1A2*, *LRRN4*, and *UPK3B*. Because there were so few (only 27) mast cells annotated in this dataset, they were not used to train the model. Lastly, only cells from the left ventricle, right ventricle, left atria, and right atria were used. The final cell type annotations used are provided in **Supplemental Table 8**. These cells form the input for downstream analyses and classifier training.

##### Development data set 1

We used the single cell RNA-seq data of Asp et al. with zone and cell type annotations to validate our random forest model^25^. Data and cell annotations were downloaded from https://www.spatialresearch.org/resources-published-da-tasets/doi-10-1016-j-cell-2019-11-025/. Cells annotated as cardiac neural crest were reannotated as immune cells based on high expression of *C1QA*, *CSF1R*, and *GIMAP4*. The final cell type annotations used are provided in **Supplemental Table 9**. In order to compute log-transformed normalized expression values, clusters were first computed (quickCluster, scran R package), followed by normalization where size factors are deconvoluted from clusters (computeSumFactors, scran), followed by log-transform normalization (logNormCounts, scater R package).

##### Development data set 2

We used the single cell RNA-seq data of Miao et al. with laterality annotations to validate our random forest model^18^. Data were downloaded from GEO (GSM4125587, GSM4125585, GSM4125586, GSM4125588). In order to compute log-transformed normalized expression values were computed as for development data set 1. Furthermore, highly variable genes, low dimensional embeddings, clustering, and cell type annotations were performed as for the unsorted hiPSC data set (see Data processing and quality control). The final cell type annotations used are provided in **Supplemental Table 10.**

##### Model fitting

To fit a model on the training set and apply it to a test data set, typically generated with different platform technology, we proceed as follows. First highly variable genes in both data sets were selected. Using the modelGeneCV2 function (scran R package) we fit the squared coefficient of variation (CV^2^) and the top 50% of genes with the largest CV^2^ and strongest deviation from the fit line were retained as highly variable genes. Additionally, genes expressed in less than 1/4^th^ of cells were filtered out. Gene passing both filters on the train and test data were scaled (for each data set independently) and kept for RF model fitting. Genes were scaled by subtracting their minimum expression value, and then dividing by their 95^th^ quartile. Next, we used the R package ranger (https://cran.r-project.org/web/packages/ranger/index.html) to derive a random forest classifier on the scaled train data (impurity importance score), using class weights to account for imbalances between cell-type labels. We then use the top genes in terms of feature importance to train a second, final random forest on the train data, which is then used to derive labels on the scaled test data set. Hyper parameters for this procedure (number of trees, number of genes for the second round of learning) were determined separately using the training data as both, test and train set, respectively. To optimize a parameter, the others were held constant while a range of values was tested and the final value was selected as the elbow point when plotting accuracy against tested parameter values. Next, the trained model is used to predict the labels on a test set. Performance is visualized using Sankey diagrams (ggplot2, https://github.com/cran/ggplot2). Cell type accuracies are calculated as the percentage of correctly classified cells. Conditional accuracies were calculated as the percentage of correctly classified cells within a given label.

##### Cell type classification

For cell type classification all cells in the training data were used in the above procedure with 300 trees and 500 important genes (**Supplementary Table 11**) as hyperparameters.

##### Anatomical zone classification

Here we focus on the anatomical zone of cardiomyocytes (CMs), and correspondingly only CMs are used in the above procedures. Hyper parameters used are: 300 trees and 100 important (**Supplementary Table 11**) genes for the second random forest.

##### Laterality classification

Here we focus on the laterality (left/right) of CMs; we proceed as discussed above, with an additional quantile normalization step after determining top-variable genes and before scaling. Hyper parameters we determined were 500 trees and 100 important genes (**Supplementary Table 11**).

##### Anomaly detection

To flag cells in the hiPSC data the model has not seen before we perform anomaly detection as follows: Cell type classification (see above) was performed and for each cell the annotated class and its class-probability were recorded. If that probability was lower than a class-dependent threshold the cell was considered an anomaly. The threshold for each class was determined as the minimum of the two 5% quantiles of probabilities of cells in the corresponding class in the two development sets.^18,25^

#### Single molecular in situ hybridization

To visualize the transcriptional expression patterns of Tnni3, Cdh5, Postn, and Wt1 in the organoids, proximity ligation in situ hybridization (PLISH) was performed as previously described with minor modifications ^44^. Briefly, the organoids were fixed with DEPC treated 4% paraformaldehyde (electron microscopy sciences, 15710S) before being embedded with OCT (Sakura, 4583). The embeded tissue were then sectioned with the thickness of 6 μm and treated with post-fix medium (3.7% formaldehyde (Sigma-Aldrich, 252549) and 0.1% DEPC (Sigma-Aldrich, D5758) for 30 min. After that, the sections were incubated with hybridization buffer (1M NaTCA, 5mM EDTA, 50mM Tris pH 7.4, 0.2mg/mL Heparin) and H probes (**supplemental Table 3**). After circulation ligation and rolling circle amplifications, the detection probes conjugated with Cy3 or Cy5 fluorophore were applied and the hybridization signal were imaged under confocal microscopy (Leica TSC SP8).

#### Immunofluorescence staining

Organoids were fixed in 4% paraformaldehyde (electron microscopy sciences, 15710S) for 1 hr. After that, the organoids were washed twice with PBS and embedded in OCT. The tissues were sectioned at 6 μm and used for staining. The immunostaining procedure was carried out as previously described ^45^. Briefly, the section slides were washed with PBS for 5 min and permeabilized with PBST (0.2% Triton X-100 in PBS) for 10 min. After that, the slides were sequentially incubated with blocking buffer (10% Goat Serum, 1% BSA, 0.1% Tween 20) for 1hr at room temperature and primary antibody in PBST with 1% BSA overnight at 4°C. The antibodies were diluted according to the manufacturer’s instructions. The mouse anti-Cardiac Troponin T (5 μg/ml, Invitrogen, MA5-12960), rabbit anti-VE-Cadherin (1:400, Cell signaling, #2500), mouse anti-Nkx2-5 (25 μg/ml, R&D Systems, #259416), rabbit anti-MYL7 (1:1000, SAB2701294) were used. The slides were further washed three times with PBS and incubated with secondary antibodies in blocking solution for 1 h at room temperature. The secondary antibodies used include goat anti-mouse 594 (10 μg/ml, A11005, Invitrogen) and goat anti-rabbit 647 (10 μg/ml, A-21245, Invitrogen). Finally, the slides were stained with DAPI for 5 min and mounted using ProLong™ Diamond Antifade Mountant (Molecular Probe, P36962). The Images were captured using Leica TCS-SP8 confocal microscope. For the quantification of the cardiac Troponin T (cTNT) expression, the mean gray values of cTNT signal were measured using Fiji ^46^ and normalized to the whole area of organoid.

#### Organoid imaging and processing

The images of beating organoids were taken under Leica DMI6000 microscope, three to ten of images were used to measure the organoid diameters. The length of both longest axis and shortest axis were measured for each organoid. Besides, the beating organoids were recorded at an interval of 50 ms with a Hamamatsu Orca-ER camera with transmitted light. The beating rates were calculated with beats/frames multiplied by frames/second.

#### Statistics

Data are presented as the mean ± standard error of the mean (SEM) for at least three replicate samples (see figure legends for additional information). Statistical significance was determined using a Student’s t-test for all quantification except RNA-seq data. Results were considered statistically significant when the P value was < 0.05 (*P < 0.05). Box plots and bar plots were generated by Prism GraphPad.

## Data Availability

Sequencing data underlying this study has been deposited at the Gene Expression Omnibus (GEO) database. Other data is available from the authors upon request.

## Code Availability

Computer code used to generate results reported in this study is available from the authors upon request.

## Author contributions

W.F. and G.L. designed the experiments; W.F. and G.L. performed the wet lab experiments and data collection; D.K, G.L, H.S and A.B designed and performed bioinformatics analysis; A.B. performed the demultiplexing, quality control and data filtering for scRNA-seq experiments; D.K and H.S. designed the label transfer procedure, H.S. implemented, trained and validated the procedure; S.J. performed the single molecular in situ hybridization and assisted in immunostaining; W.F., H.S., A.B, D.K. and G.L. performed data analysis; W.F., H.S., A.B., D.K., and G.L. prepared the manuscript.

## Acknowledgement

The authors acknowledge the Allen Cell Collection and Joseph C. Wu, MD, PhD at the Stanford Cardiovascular Institute for kindly providing us the WTC (ACTN2-eGFP) and SCVI114 iPSC line, respectively. We thank David M. Patterson and Christopher S. McGinnis from Dr. Zev J. Gartner lab for their kind supply of the lipid based barcoding reagents and suggestions on the experimental steps and data analysis. We also would like to thank Timothy Feinstein from the Department of Developmental Biology, University of Pittsburgh school of medicine, for the help in cell imaging process and valuable discussions. Finally, we thank Josh Michel from Rangos Flow Cytometry Core, Children’s Hospital of Pittsburgh, for his assistance with cell sorting. This work was founded by the National Institutes of Health (HL133472).

## Footnotes

1 We use “cell” as a shorthand for cell/10X-barcode in an abuse of notation, since multiplets are not single cells by definition.

